# Ubiquitin Substrates Dramatically Increase Ataxin3 Deubiquitinating Activity: Allosteric crosstalk connects three distinct sites

**DOI:** 10.1101/2020.05.05.078998

**Authors:** Maya V. Rao, Kimberly C. Grasty, Prajakta D. Mehetre, Patrick J. Loll

**Affiliations:** Department of Biochemistry & Molecular Biology, Drexel University College of Medicine, Philadelphia, PA 19102 USA; UCB Biosciences Inc., Bedford MA 01730 USA

## Abstract

Ataxin3 is the founding member of the MJD family of deubiquitinating enzymes, and plays important roles in maintaining protein homeostasis and promoting DNA repair. The enzyme also contains a polyglutamine tract of variable length, and in its expanded form the protein becomes the causative agent of a neurodegenerative disorder known as Machado-Joseph disease. *In vitro,* ataxin3 displays low catalytic activity, prompting questions about how the enzyme is regulated and what signals might lead to its activation. Recently, it has been demonstrated that ataxin3 activity can be stimulated by either mono-ubiquitination or high concentrations of free ubiquitin. Here, we show that ubiquitin conjugates with cleavable bonds can stimulate ataxin3 activity much more strongly than free ubiquitin, with physiological levels of these conjugates increasing activity up to 60-fold over basal levels. Our data are consistent with a model in which ubiquitin conjugates activate the enzyme allosterically by binding in a site adjacent to the catalytic center, known as Site 1. We further show that two additional ubiquitin-binding sites in the enzyme work in concert to modulate enzyme activation, and we propose a model in which ubiquitin conjugates bridge these two sites to drive the enzyme into a high-activity conformation.

**Significance:** Ubiquitin signaling networks modulate almost all aspects of eukaryotic biology, and their outputs reflect the dynamic balance between ubiquitin attachment and removal. The latter process is catalyzed by deubiquitinating enzymes (DUBs), which must be carefully regulated to ensure that their activities are applied appropriately. Ataxin3 is a DUB that participates in quality-control pathways that support cellular health; however, the regulation of its activity has remained poorly understood. Here, we show that ataxin3 can be dramatically activated by naturally occurring ubiquitin species, and that this activation involves a previously uncharacterized interplay between three distinct sites on the enzyme. Our improved understanding of ataxin3 regulation provides insights into allosteric mechanisms that may prove applicable to other enzymes in the ubiquitin universe.

## Introduction

Ubiquitin signaling affects almost all aspects of eukaryotic biology, including protein homeostasis, DNA repair, cell cycle, vesicular trafficking, receptor-mediated signaling, and transcription (1-6). The addition of ubiquitin signals is balanced by their removal, which is catalyzed by a wide array of deubiquitinating enzymes (DUBs): ubiquitin-specific peptidases that edit or remove ubiquitin modifications (7). Ubiquitin linkages can adopt many different topologies (8), and consistent with the diversity of possible substrates, many different DUBs exist; for example, the human genome encodes roughly 100 such enzymes, divided into seven distinct families (7). To avoid indiscriminate or inappropriate DUB processing, cells have evolved multiple layers of regulatory control (9, 10). These include regulation of the abundance and localization of different DUBs (11, 12), as well as mechanisms for direct control of DUB activities. Examples of the latter include substrate-assisted catalysis (13, 14); post-translational modifications such as phosphorylation, ubiquitination, or SUMOylation (15); control of the redox state of active-site cysteines (16); and allosteric mechanisms (17, 18).

The smallest of the seven DUB families is the MJD, or Josephin, family. Four family members are known in humans, the best-studied of which is ataxin3, which first came to prominence by virtue of its status as a polyglutamine protein (19, 20). Mutations that expand the ataxin3 polyglutamine tract lead to the progressive neurological disorder known as <u>M</u>achado-<u>J</u>oseph <u>d</u>isease (MJD) (21). However, while mutant ataxin3 is a disease-causing agent, wild-type ataxin3 plays important cytoprotective roles in normal cellular physiology, supporting protein quality-control pathways and DNA repair (22-28).

Different splice variants of ataxin3 are known, but each contains approximately 350 residues (29). The N-terminal half of the protein forms a catalytic “Josephin” domain, which is common to all MJD family members and adopts a papain-like cysteine protease fold (30-33). The ataxin3 Josephin domain contains two different ubiquitin-binding sites (34). The first (known as Site 1) binds and positions the distal ubiquitin partner in the scissile bond; the second (Site 2) lies on the opposite side of the Josephin domain, and may play a role in positioning poly-ubiquitin substrates. In contrast to the ordered Josephin domain, the C-terminal half of ataxin3 forms an intrinsically disordered domain (35, 36); this region contains a nuclear localization sequence (37), a p97/VCP-binding motif (38), the polyglutamine tract, and a third ubiquitin-binding site, formed by two tandem <u>u</u>biquitin-interacting <u>m</u>otifs (UIMs; Figs. 1*A* and *B*) (39).

**Figure 1.**
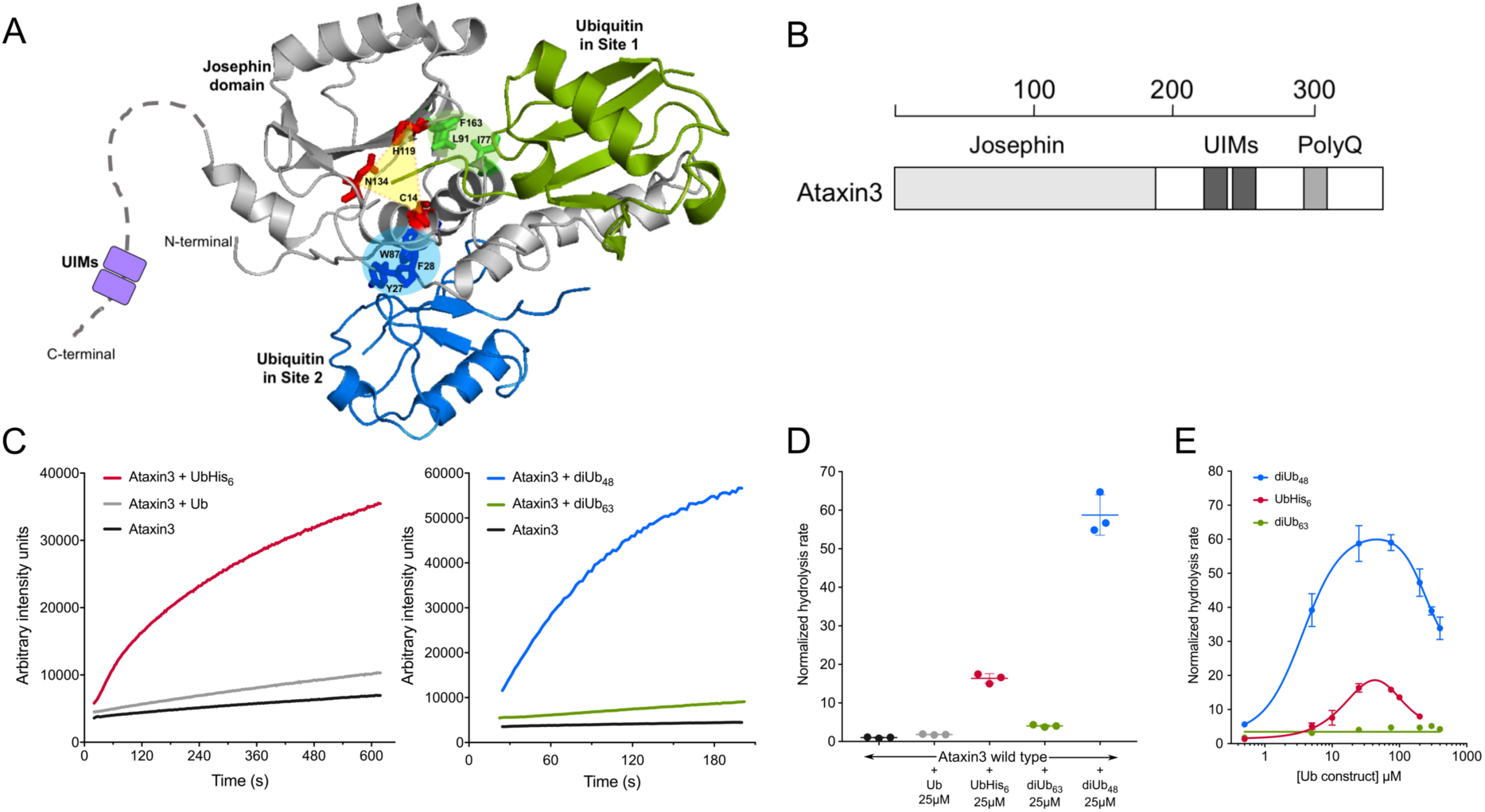
Mono- and di-ubiquitin stimulate ataxin3’s DUB activity. *(A)* Three-dimensional architecture of ataxin3 and its ubiquitin-binding sites (coordinates from PDB entry 2JRI). The structure of the Josephin domain is shown in gray. The residues of the catalytic triad are shown in red stick representation, connected by a yellow triangle representing the catalytic center. Ubiquitin molecules are shown bound in Site 1 (green) and Site 2 (blue); key ataxin3 residues controlling ubiquitin recognition in each site are shown (Site 1: I77, L91, & F163; Site 2: Y27, F28, and W87). The disordered ataxin3 C-terminal domain is represented by a dashed line, with the position of the tandem UIMs being indicated by a pair of purple boxes. *(B*) Schematic view of the ataxin3 protein. Scale bar at top shows length in amino acids. *(C)* Representative raw fluorescence traces showing stimulation of ataxin3 hydrolysis activity by 25 µM ubiquitin constructs, as measured by the Ub-AMC cleavage assay. The left panel shows stimulation by ubiquitin (Ub; gray) and ubiquitin-His_6_ (UbHis_6_; red), and the right panel shows stimulation by K48-linked di-ubiquitin (diUb_48_; blue) and K63-linked di-ubiquitin (diUb_63_; green). Note that the y-axis scales differ in the two panels. In both panels, basal activity of ataxin3 alone is shown in black. *(D)* Initial rates of Ub-AMC hydrolysis for ataxin3 ± 25 µM ubiquitin and ubiquitin conjugates. Rates are normalized to the basal (unstimulated) activity for ataxin3. *(E)* Dose response for ataxin3 stimulation by three ubiquitin conjugates. Error bars in *(D)* & *(E)* show standard deviations calculated for three or more replicates.

Ataxin3 cleaves long ubiquitin chains more efficiently than smaller substrates, and is selective for K63-linked linear chains and K48/K63-linked branched chains (23, 33, 40, 41). However, its overall activity level is markedly lower than that of many other DUBs, suggesting that activation of catalysis might be a key aspect of the enzyme’s regulation. Recently ataxin3 has been shown to be monoubiquitinated on Lys-117, leading to an increase in catalytic activity (40, 42). This activation has been suggested to occur via partitioning of the tethered ubiquitin molecule into Site 1, thereby driving the enzyme toward a more active conformational state (43). This model predicts that free ubiquitin binding in Site 1 should activate in a similar manner, and indeed, free ubiquitin has been demonstrated to activate the enzyme, although high concentrations are required. Following up on this observation, we have compared the ability of free ubiquitin and small ubiquitin conjugates to activate ataxin3. We show that ubiquitin conjugates bearing cleavable groups at the C-terminus can stimulate ataxin3 activity much more strongly than ubiquitin itself, and that multiple sites on the enzyme contribute to modulating this allosteric activation.

## Results

### Mono- and di-ubiquitin stimulate ataxin3’s deubiquitinating activity

There is increasing evidence that both ubiquitin ligases and deubiquitinating enzymes can be regulated by the noncovalent binding of ubiquitin itself (9, 44). In particular, the enzymatic activity of the deubiquitinating enzyme ataxin3 has been reported to be stimulated by free ubiquitin (43), and we chose to examine the basis of this stimulation.

To monitor ataxin3 activity, we used the synthetic substrate ubiquitin-7-amino-4-methoxycoumarin (Ub-AMC), which consists of the fluorophore AMC conjugated to the C-terminal of ubiquitin by an amide linkage (45). When attached to ubiquitin, AMC is only weakly fluorescent, but once cleaved, its fluorescence increases substantially; the Ub-AMC substrate therefore affords a sensitive and continuous method for monitoring deubiquitinating activity. We first measured the activity of purified wild-type ataxin3 alone under steady-state conditions, using 20 nM ataxin3 with 1 µM Ub-AMC substrate (Figs. 1*C* and *D*). The initial rate was derived and all subsequent activity measurements were normalized to this basal activity level (Fig. S1; the basal activity level is represented by a dotted line in Figs. 2 through 4).

**Figure 2.**
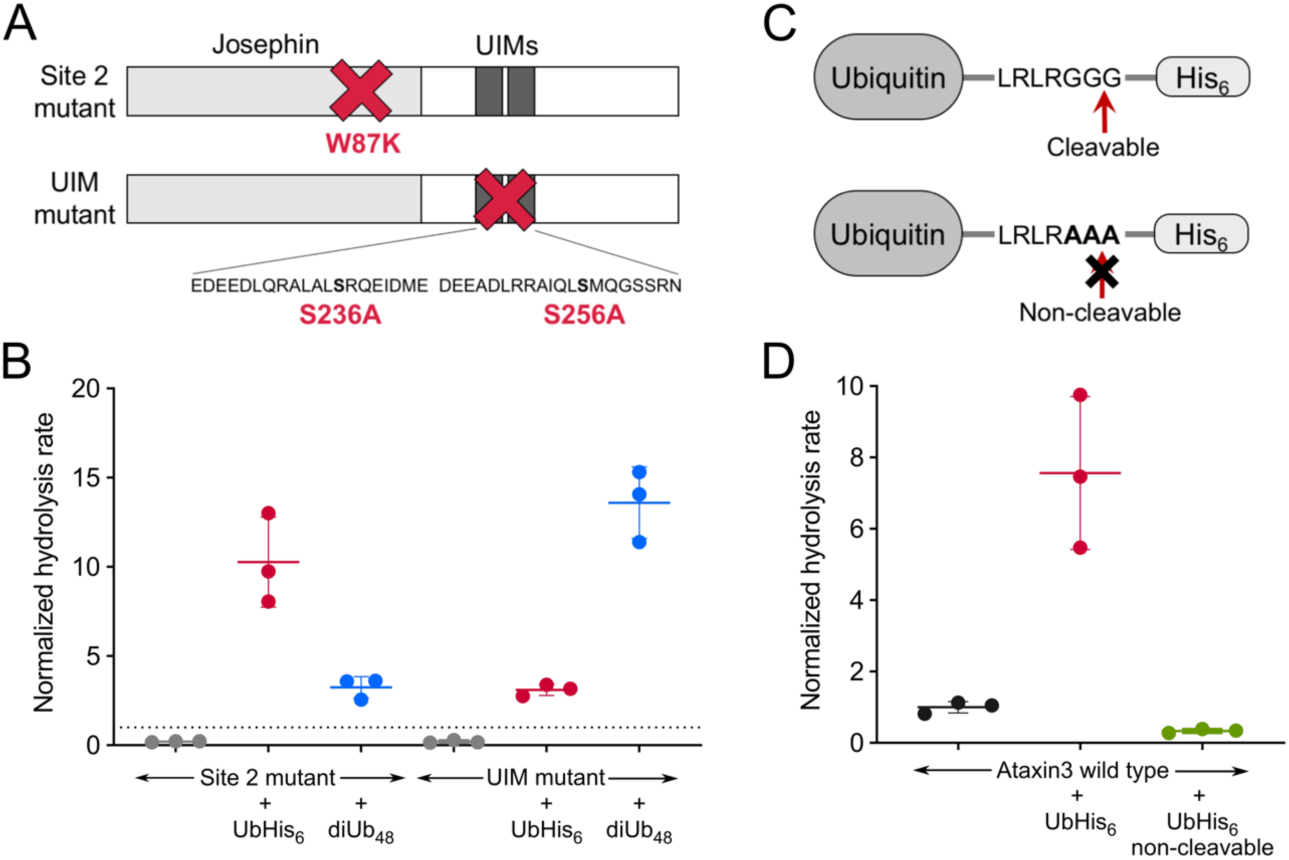
Using single-site mutations to probe ubiquitin activation of ataxin3. *(A)* Schematic views of the two single-site ataxin3 mutants. The Site-2 mutant carries the W87K mutation, while the tandem UIM mutant carries both the S236A and S256A mutations. *(B)* Effect of 25 µM concentrations of different ubiquitin conjugates on the two single-site ataxin3 mutants. *(C)* Schematic view of the UbHis_6_ conjugate in cleavable and non-cleavable forms. DUBs cleave ubiquitin conjugates immediately after Gly-76, as shown by the red arrow; the non-cleavable analog replaces Gly-75, Gly-76, and the following residue with alanines. *(D)* Effect of 10 µM concentrations of UbHis_6_ conjugates on ataxin3 activity. All rates shown (*n* ≥ 3) are normalized to the basal rate for wild-type ataxin3 (dotted line).

We then tested the effects of various monomeric and dimeric ubiquitin constructs on ataxin3’s deubiquitinating activity, using an excess (25 µM) of either native ubiquitin (Ub), C-terminally His_6_-tagged ubiquitin (UbHis_6_), K48-linked di-ubiquitin (diUb_48_), or K63-linked di-ubiquitin (diUb_63_). We note that three of these constructs—UbHis_6_, diUb_48_, and diUb_63_—can function as substrates for ataxin3, but their cleavage will not be detected in the Ub-AMC assay. All four of the ubiquitin constructs tested stimulated Ub-AMC hydrolysis (Figs. 1*C* and *D*), with the most potent construct, diUb_48_, increasing enzymatic activity by approximately 60-fold. The other three constructs provided lower levels of stimulation, ranging from approximately two-fold for Ub to 18-fold for UbHis_6_. The concentration dependence of this stimulatory effect was measured, and for the two most potent constructs (diUb_48_ and UbHis_6_) the dose-response was observed to be bimodal, with maximal stimulation occurring in the concentration range 25 – 75 µM, above which enzyme activity decreased (Fig. 1*E*).

### Identification of the site on ataxin3 that mediates ubiquitin stimulation

Reasoning that ubiquitin must bind ataxin3 in order to stimulate it, we set out to identify the binding site (or sites) responsible. Three ubiquitin-binding sites are known on ataxin3, namely the catalytic site, referred to as Site 1; a second site centered around Y27, F28, and W87 on the Josephin domain, known as Site 2 (34); and the tandem UIMs, which lie between residues 224 and 263 in the C-terminal region of the enzyme (Fig. 1*A*) (39, 46). Site 1, being the catalytic site, cannot be altered without compromising enzyme activity (34), which precludes mutagenesis within this region. We therefore introduced mutations into the other two sites and studied their effects on ataxin3 stimulation. Site 2 and the tandem UIMs were rendered incapable of binding ubiquitin by the W87K mutation and the paired S236A/S256A mutations, respectively (Fig. 2*A*) (34, 47). We found that both the Site-2 mutant and the UIM mutant were stimulated by both UbHis_6_ and diUb_48_, albeit to differing extents (Fig. 2*B*), and therefore conclude that neither Site 2 nor the UIM is absolutely required for ubiquitin to stimulate ataxin3 activity. Accordingly, the stimulatory effect of ubiquitin appears to be mediated through Site 1, as suggested by Faggiano *et al.* (43).

Activation via ubiquitin binding in Site 1 would explain the bimodal dose-response activation curves observed for UbHis_6_ and diUb_48_ (Fig. 1*E*). The enzymatic assay used reports only production of AMC, which requires that Ub-AMC binds in Site 1; however, if the stimulatory ubiquitin species also binds in this site, it will compete with Ub-AMC, and at sufficiently high concentrations competition will outweigh stimulation and thus reduce the rate of Ub-AMC hydrolysis. We tested this idea by using a non-hydrolyzable analog of UbHis_6_ in which the Gly-Gly-X DUB cleavage site was mutated to Ala-Ala-X (Fig. 2*C*). We reasoned that if this molecule bound in Site 1, it would not be turned over, and should therefore inhibit, rather than stimulate. Indeed, at 10 µM, the non-hydrolyzable analog reduces the Ub-AMC hydrolysis rate to roughly one-third the basal level, whereas the same concentration of UbHis_6_ stimulates ataxin3 activity eight-fold above basal (Fig. 2*D*). Taken together, these results support the notion that mono- and di-ubiquitin constructs stimulate ataxin3 activity by binding in Site 1.

### Behavior of the Site-2 + UIM double-site mutant

Neither single-site mutant reached full wild-type activity, even in the presence of high concentrations of UbHis_6_ or diUb_48_ (Fig. 2*B*). Since Site 1 is intact in both single-site mutants, this result suggests that Site-1 occupancy is not the only factor affecting ataxin3 catalysis. To assess whether Site 2 and the UIMs also help modulate enzyme activity, we constructed an ataxin3 “double-site mutant” in which both Site 2 and the tandem UIMs were mutated (W87K/S236A/S256A) (Fig. 3*A*). This double-site mutant enzyme behaved differently than the wild-type and single-site mutant proteins, in ways that we had not predicted. For example, UbHis_6_ stimulated the wild-type and double-site mutant enzymes to similar degrees, whereas diUb_48_ stimulated unequally: 25 µM diUb_48_ stimulated the wild-type enzyme ∼60-fold, but increased the activity of the double-site mutant by only four-fold (Figs. 1*D* and 3*B*). Further, the basal (unstimulated) activity of the double-site mutant was higher than that of the wild-type enzyme, while basal activities of both single-site mutants were lower than wild type (Figs. 2*B* and S1). The latter observation suggested that Site 2 and the UIM-containing C-terminus might be working in concert to modulate enzyme activity. Accordingly, we examined the activity of the isolated Josephin domain, which contains Site 2 but lacks the entire C-terminal region, including the tandem UIMs. For this reason, we hypothesized the Josephin domain would behave similarly to the UIM mutant of the full-length enzyme.

**Figure 3.**
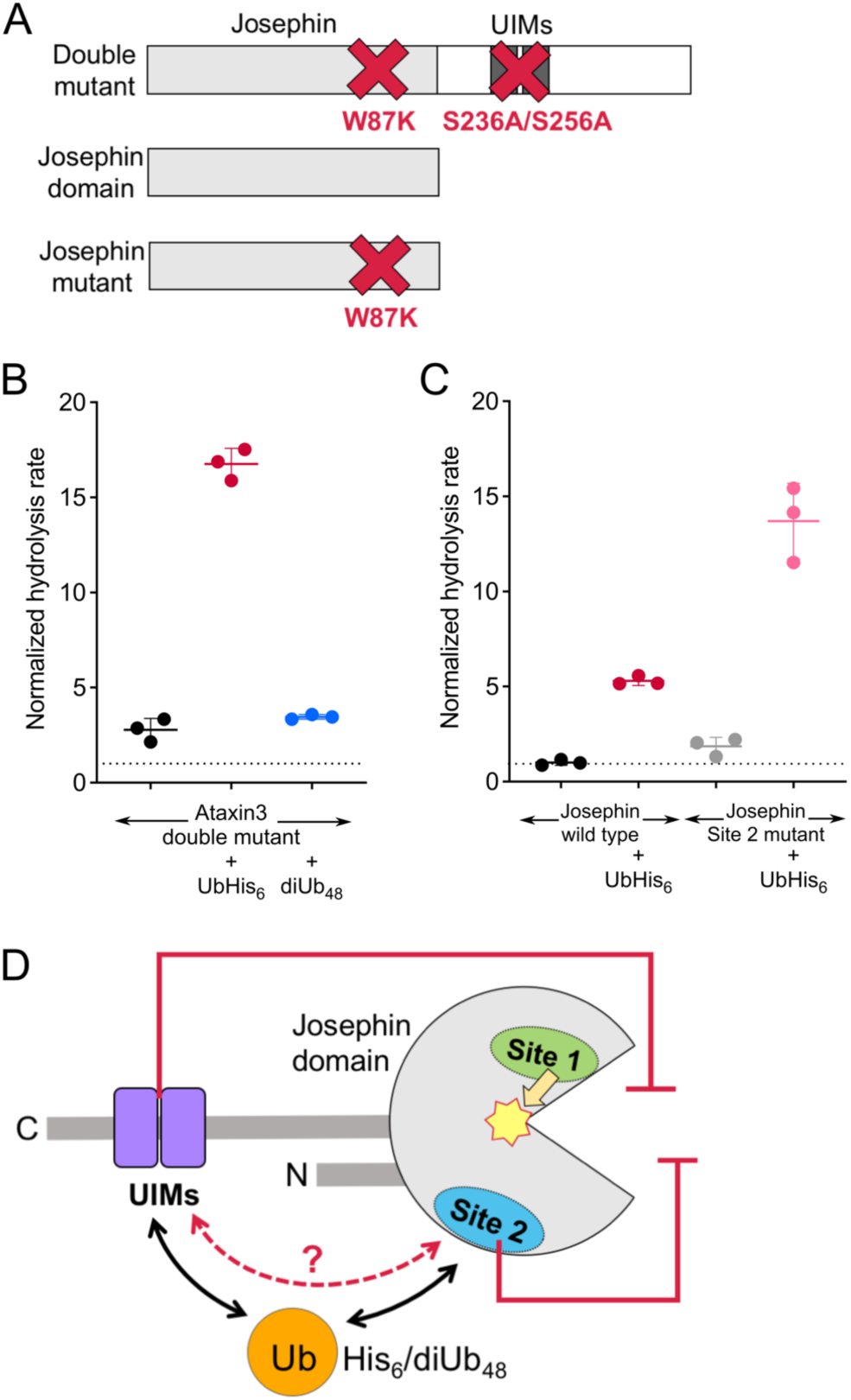
Analysis of an ataxin3 double-site mutant suggests synergy between Site 2 and the tandem UIMs. *(A)* Schematic views of the ataxin3 double-site mutant (bearing the W87K/S236A/S256A mutations), the wild-type Josephin domain, and its Site-2 mutant form (bearing the W87K mutation). *(B)* Basal and stimulated activities of the ataxin3 double-site mutant ± 25 µM UbHis_6_ and diUb_48_. *(C)* The activities of wild-type and Site-2 mutant forms of the Josephin domain ± 75 µM UbHis_6_. *(D)* Synopsis of ubiquitin effects on ataxin3, as inferred from mutagenesis experiments: Binding of ubiquitin species on Site 1 provides a stimulatory effect on the catalytic center; binding of ubiquitin species at *either* Site 2 or the tandem UIMs inhibits activity; simultaneous binding of ubiquitin species in *both* Site 2 and the tandem UIMs relieves this inhibitory effect. Average rates (*n* ≥ 3) are normalized to the basal rates for wild-type full-length ataxin3 (panel *C*) or the wild-type Josephin domain (panel *D*). Reference rates are indicated by dotted lines.

### Stimulation of activity in the Josephin domain recapitulates the behavior of full-length ataxin3

To determine how the isolated Josephin domain responds to ubiquitin stimulation, we expressed and purified a construct containing residues 1-190 and used it in Ub-AMC activity assays (Fig. 3*A*). As has been noted previously (33), the activity of the Josephin domain is substantially lower than that of full-length ataxin3 (Fig. S1). In fact, in order to reliably measure the Josephin domain’s basal activity it was necessary to alter the assay protocol, using higher enzyme and substrate concentrations than those used with the full-length protein (see SI Methods & Table S2).

Since the Josephin domain lacks the entire C-terminal sequence of the protein, it can be regarded as roughly equivalent to the UIM mutant of the full-length protein, insofar as both constructs lack any functional UIMs. We found that the Josephin domain could be stimulated by UbHis_6_, but it could never reach the activity levels seen with the full-length protein (Fig. 3*C*). Thus, its behavior is consistent with that of the UIM mutant of the full-length protein. Similarly, the Josephin domain bearing the Site-2 W87K mutation is roughly equivalent to the double-site mutant of the full-length protein, in that it contains neither the tandem UIMs nor a functional Site 2. The Josephin Site-2 mutant demonstrated a higher basal activity than the wild-type Josephin, and addition of UbHis_6_ increased its activity to higher levels than it does for the wild-type Josephin (Fig. 3*C*). Hence the behavior of this mutant is consistent with that of the double-site mutant of the full-length protein. Therefore, at least in broad strokes, the modulation of Josephin domain activity by UbHis_6_ is comparable to that seen in the full-length enzyme.

### A model for the interplay of Site 2 and the tandem UIMs

At high ubiquitin concentrations, neither of the single-site mutants of the full-length protein can reach full wild-type levels of activity. However, when these two mutants are combined, the resulting double-site mutant can be fully stimulated, to levels equivalent to those observed for the wild-type protein. It therefore appears that when ubiquitin binds *only* to Site 2 or *only* to the tandem UIMs, it somehow exerts an inhibitory effect on the enzyme and/or fails to fully activate; however, when ubiquitin binds simultaneously to *both* sites, this inhibition is relieved and/or full activation is achieved (Fig. 3*D*). This model predicts different behaviors for the wild-type, single-site mutant, and double-site mutant proteins that are consistent with our observations. For the wild-type protein, ubiquitin conjugates bind in Site 1, which stimulates the enzyme, and also bind to both Site 2 and to the tandem UIMs, thus cancelling out any inhibitory effects. For the single-site mutants, stimulation from Site 1 will be partially offset by inhibition caused by binding in either Site 2 or the UIM, resulting in sub-maximal activity levels. For the double-site mutant, however, neither Site 2 nor the UIM is occupied, so there is no inhibition and the enzyme is fully stimulated.

### Evidence for interaction between Site 2 and the tandem UIMs

The model described above postulates that enzyme activity is sensitive to the occupancies of both Site 2 and the tandem UIMs, raising the possibility that these two sites are communicating with one another. To probe this possibility, we first asked whether the two sites function in *cis* or in *trans*. We prepared a chimeric protein in which the wild-type tandem UIM sequence was fused to the C-terminus of <u>m</u>altose-<u>b</u>inding <u>p</u>rotein (MBP), and then tested whether this MBP-UIM fusion could restore full activation of the ataxin3 UIM mutant in the presence of high concentrations of UbHis_6_. We observed that neither the MBP-UIM construct nor an MBP control protein had any effect on the activity of the UIM mutant (Fig. 4*A*). Hence, Site 2 and the tandem UIMs must be present in *cis* (on the same polypeptide chain) in order to affect enzyme activity.

**Figure 4.**
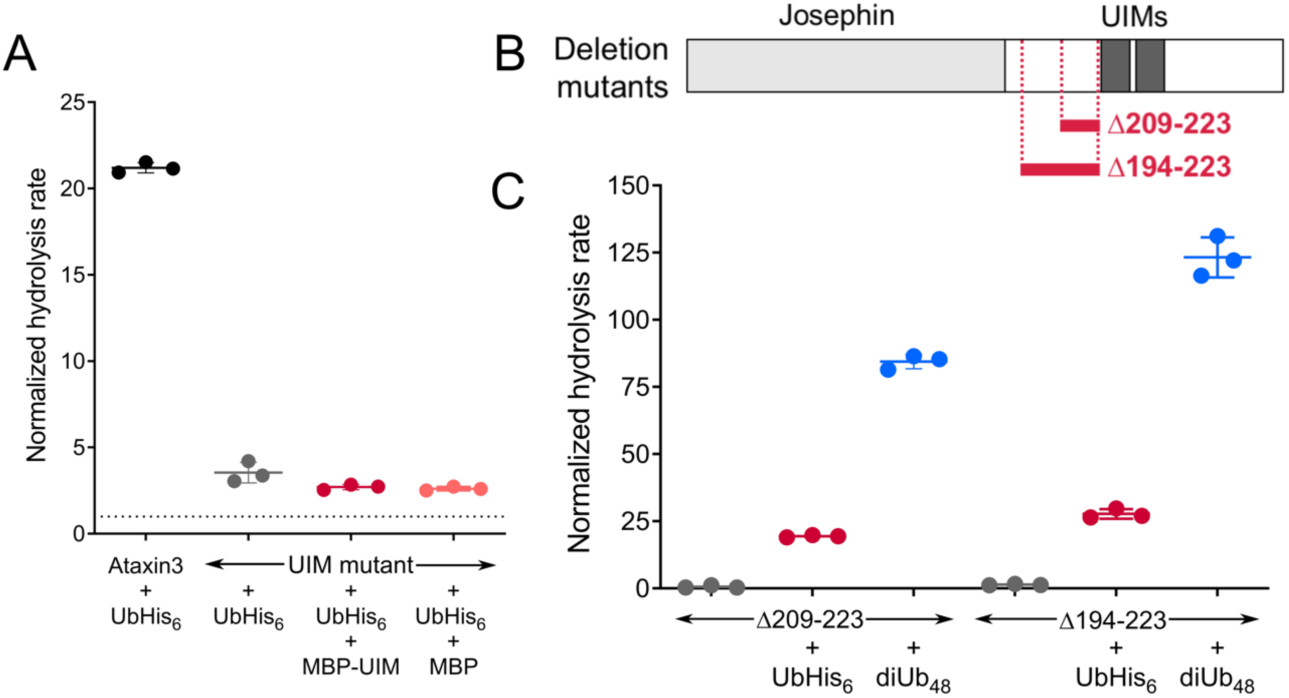
Site 2 and the UIMs communicate intramolecularly, in a proximity-dependent manner. *(A)* The tandem UIMs must be present *in cis* to affect ataxin3 DUB activity. Shown are normalized activities for wild-type ataxin3 or the tandem UIM mutant in the presence of 25 µM UbHis_6_. The ataxin3 UIM mutant shows greatly reduced stimulation by UbHis_6_; adding back a functional copy of the tandem UIMs *in trans* (as a maltose-binding protein fusion; MBP-UIM) fails to restore stimulation. In a control experiment, addition of MBP alone has no effect on activity. MBP and the MBP-UIM fusion were both used at a concentration of 1 µM. *(B)* Schematic views of the ataxin3 deletion mutants. In-frame deletions of fifteen residues (Δ209-223) or thirty residues (Δ194-223) were made immediately upstream of the UIMs. *(C)* The activities of the truncated proteins ± 25 µM UbHis_6_ or diUb_48_; reducing the linker length separating the Josephin domain from the tandem UIMs enhances stimulation by both UbHis_6_ and diUb_48_. Average rates (*n* ≥ 3) are normalized to the basal rate for wild-type ataxin3 (shown as a dotted line in panel *A*).

The requirement that the two sites be present on the same polypeptide chain suggests that the tandem UIMs may be coming into close proximity with Site 2. Alternatively, the specific amino-acid sequences flanking the UIM may contribute to the modulation of enzyme activity. To address these possibilities, we constructed two deletion mutants that shorten the linker connecting the tandem UIMs to the Josephin domain. We excised either 15 or 30 residues immediately upstream of the UIM, producing constructs Δ209-223 and Δ194-223, respectively (Fig. 4*B*). These deletion mutants proved to have higher basal activities than wild-type ataxin3, and stimulation of the mutants by either UbHis_6_ or diUb_48_ resulted in higher activity levels than could be achieved with the wild type (Fig. 4*C*). Both basal and stimulated activity levels were higher for the larger deletion (Δ194-223); in the presence of 25 µM diUb_48_, this construct achieved an activity level twice that of wild-type ataxin3, which represents a greater-than 100-fold stimulation relative to the basal wild-type activity. Hence, linker length, rather than the specific linker sequence adjoining the UIMs, is the more important variable in the interplay between the tandem UIMs and the Josephin domain, bolstering the notion that the two sites must come into proximity.

To determine whether activation by ubiquitin is associated with a change in distance between Site 2 and the tandem UIMs, we constructed a fluorescent reporter construct for ataxin3. In this construct, YPet was fused to the N-terminus of the enzyme and CyPet was fused immediately downstream of the tandem UIMs (Fig. 5*A*) (48). The N-terminus of ataxin3 lies close to Site 2; in contrast, the tandem UIMs are separated from the Josephin domain by approximately 85 residues, which are believed to be largely disordered (35). Hence, the CyPet partner is expected to occupy a wide range of different positions, many of them far from the Josephin domain. If activation by ubiquitin is accompanied by a conformational change that brings Site 2 into close proximity with the tandem UIMs, this should bring the CyPet much closer to the Josephin domain and increase FRET efficiency for the YPet-CyPet pair. We first confirmed that the enzymatic activity of the fluorescent reporter was comparable to that of the wild-type enzyme, and that the reporter could be stimulated by UbHis_6_ and diUb_48_ proteins in a manner similar to that seen for the wild-type (Fig. 5*B*). We then examined the effect of UbHis_6_ and diUb_48_ on the FRET efficiency of the reporter construct. Both of the ubiquitin proteins elicited significant changes in the fluorescent spectrum of the reporter, consistent with increased levels of FRET (Fig. 5*C*). As seen for enzyme activation, diUb_48_ proved to be more potent than UbHis_6_, and in fact the FRET dose-response curves for UbHis_6_ and diUb_4_ closely mirror the ascending legs of their enzyme-activity dose-response curves (compare Figs. 5*D* and 1*E*). We performed control experiments with SUMO, a protein similar in size to but distinct from ubiquitin, and found no nonspecific effects (e.g., crowding) on the FRET efficiency of the YPet-ataxin3-CyPet construct (Fig. S3*A*). Further controls showed that neither UbHis_6_ or diUb_48_ altered the fluorescence behavior of either YPet or the ataxin3-CyPet proteins (Fig. S3*B*). Hence, the FRET data provide strong evidence for a model in which ubiquitin drives a conformational change in ataxin3 that juxtaposes Site 2 with the tandem UIMs (Fig. 5*E*).

**Figure 5.**
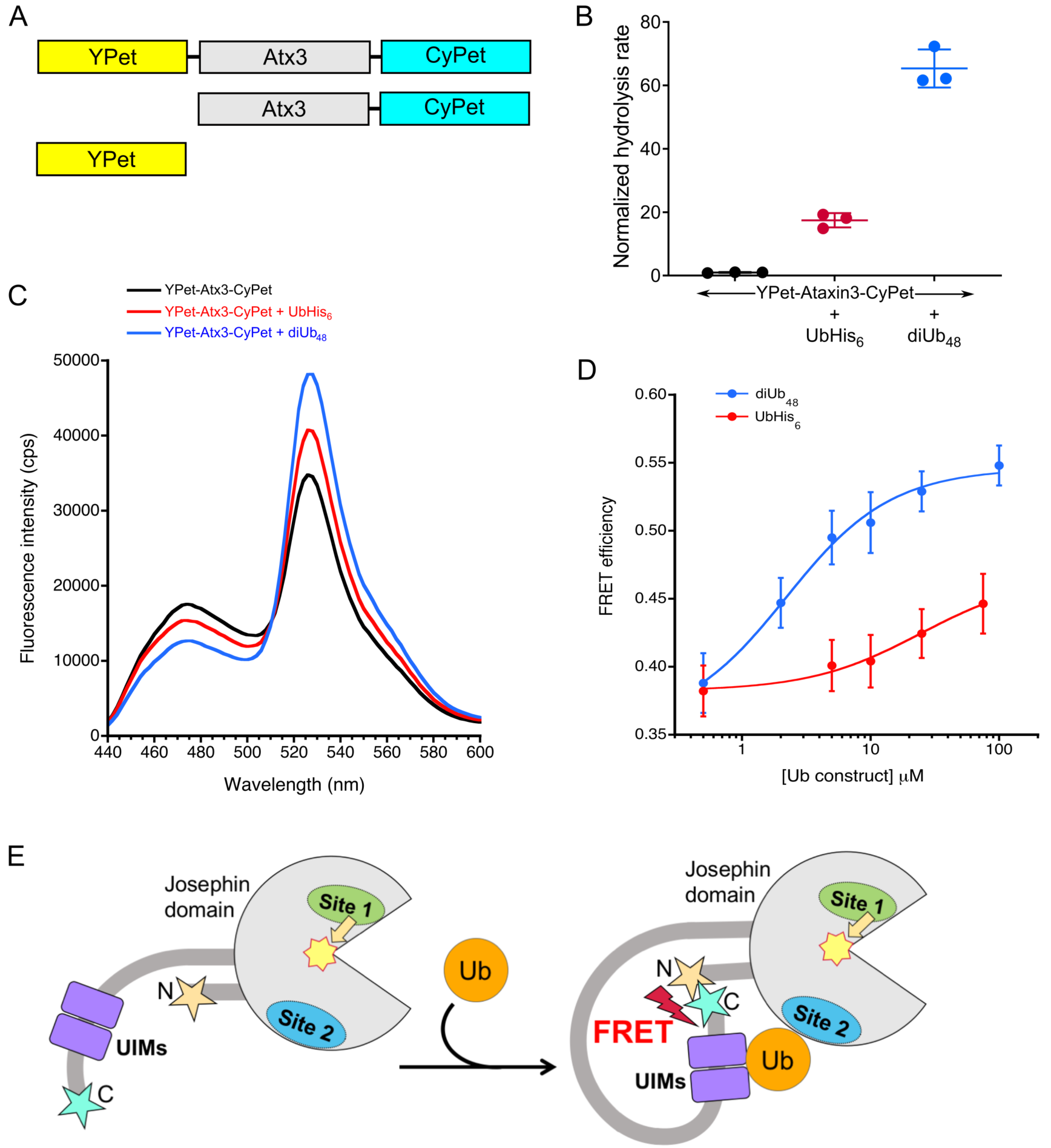
FRET evidence for communication between Site 2 and the tandem UIMs. *(A)* Schematic representation of the YPet-ataxin3-CyPet FRET sensor, as well as the YPet and ataxin3-CyPet control proteins. *(B)* The YPet-ataxin3-CyPet fusion protein is enzymatically active and is activated normally by 25 µM concentrations of UbHis_6_ and diUb_48_. Average rates (*n* = 3) are normalized to the basal activity rate for the YPet-ataxin3-CyPet protein. *(C)* Ubiquitin conjugates enhance FRET in the CyPet/YPet donor-acceptor pair. Shown are representative fluorescence spectra for YPet-ataxin3-CyPet, either alone (black) or in the presence of 25 µM UbHis_6_ and diUb_48_ (red and blue, respectively). This experiment was repeated 3-5 times with comparable results. No FRET is observed for equimolar mixtures of YPet and ataxin3-CyPet, either in the presence or absence of ubiquitin conjugates (Fig. S3*B*). *(D)* Dose-response for stimulation of FRET efficiency in the YPet-ataxin3-CyPet sensor (*n* ≥ 5). *(E)* Model for how ubiquitin binding brings together Site 2 and the tandem UIMs, enhancing FRET efficiency in the process.

## Discussion

Ataxin3’s DUB activity contributes to maintaining protein homeostasis under both normal and stressed conditions, and therefore the regulation of this activity has significant consequences for cellular health. One likely form of ataxin3 regulation is substrate-assisted catalysis, as manifested by the enzyme’s selectivity for long ubiquitin chains and certain branched species (33, 41, 49). In addition, the enzyme is regulated via monoubiquitination at Lys-117, which leads to a modest four-to-six-fold increase in activity (40, 42, 43). The K117-linked ubiquitin molecule stimulates ataxin3 activity by partitioning into Site 1, which promotes a catalytically active conformation of the enzyme (43). Presumably, the enzyme remains in this active conformation for some period of time after the K117-ubiquitin leaves, and is therefore primed to accept and hydrolyze substrate. This model predicts that binding of free ubiquitin in Site 1 should also stimulate enzyme activity, and indeed this is observed. However, unlike a ubiquitin molecule covalently tethered at Lys-117, free ubiquitin enjoys no proximity advantage that would increase its local concentration around Site 1. As a result, unphysiologically high concentrations of free ubiquitin (∼1-2 mM) are required to produce significant stimulation of enzyme activity (43).

We now show that ubiquitin constructs other than monomeric ubiquitin also stimulate ataxin3 activity, and can be much more potent activators of the enzyme. In particular, ubiquitin conjugates that contain cleavable bonds at their C-termini (*e.g.* UbHis_6_ or di-ubiquitin) are strongly stimulating, which may indicate that binding of a *bona fide* substrate is required to properly organize the enzyme’s active site. DiUb_48_, the most potent construct we tested, stimulates ataxin3’s activity by roughly 60-fold at a concentration of 25 µM. Since the total intracellular level of ubiquitin (free + conjugated) is estimated to be 10-20 µM (50), it is plausible that ubiquitin conjugates could achieve sufficiently high local concentrations near ataxin3 to strongly activate the enzyme, arguing for the physiological relevance of this effect. The stimulation caused by diUb_48_ is roughly an order of magnitude greater than that provided by mono-ubiquitination at Lys-117; however, the effects of ubiquitination at Lys-117 could possibly synergize with the effects of ubiquitin conjugates binding noncovalently to Site 2 and the tandem UIMs.

Our single-site mutant data reveal that neither Site 2 nor the tandem UIMs are absolutely required for stimulation, which is consistent with the model outlined by Faggiano *et al.* (43), in which ubiquitin stimulates by binding to Site 1. A counter-intuitive aspect of this model is stimulation by competitors—i.e., one ubiquitin species can stimulate cleavage of a different species, even though both compete to bind in the active site. This model is reminiscent of the mechanism of glucokinase activation, in which substrate binding in the active site promotes a conformational change that persists even after the product diffuses away (51, 52). The details of how ubiquitin binding in Site 1 promotes activity remain unknown; however, since ataxin3 is known to undergo significant structural fluctuations in the absence of substrate (53), the activating effects of ubiquitin may involve narrowing the enzyme’s conformational ensemble.

While our mutational analysis supports the Site-1 stimulation model, at the same time it indicates that ataxin3 regulation is not limited to Site 1, but also extends to Site 2 and the tandem UIMs. Occupancy of either individual site by ubiquitin appears to exert an inhibitory effect on ataxin3 activity, while simultaneous occupation of both sites relieves this inhibition (and/or provides additional stimulation). Ubiquitin binding at Site 2 might affect catalysis by perturbing the conformation of the Josephin domain; however, it is less straightforward to rationalize how ubiquitin binding to the tandem UIMs could have a similar effect, given the UIMs’ distance from the catalytic center. However, we note that expansion of the ataxin3 polyglutamine tract—which is adjacent to the UIMs—promotes local unfolding within the Josephin domain (54, 55), proving that information about the protein’s C-terminal region is somehow communicated with the catalytic domain.

Mutations in Site 2 and the tandem UIMs are not additive when combined in a double-site mutant, providing further support for inter-site communication. In order to communicate, the two sites must be present *in cis*, and the length of the connecting sequence controls how strongly they affect ataxin3 activity. Together, these observations suggest that Site 2 and the tandem UIMs physically approach one another. We tested this notion using a FRET reporter construct, and confirmed that ubiquitin conjugates reduce the distance between ataxin3’s N- and C-termini. Addition of ubiquitin conjugates increases the FRET efficiency for the CyPet-YPet pair from ∼0.37 to a maximum value of ∼0.55; assuming a Forster distance of 49 Å for the FRET pair (56), this corresponds to a decrease in mean fluorophore separation from 54 Å to 47 Å. While detailed interpretation of these values is impossible in the absence of high-resolution structural information, we note that the 54 Å value obtained in the absence of any ubiquitin conjugates is lower than would be predicted from a purely random-coil model for the ataxin3 C-terminal domain (Supplemental Information & Fig. S4). The reasons for this discrepancy are not obvious; however, what is clear that is addition of various ubiquitin species significantly reduces this distance, consistent with a ubiquitin-triggered conformational change that moves the tandem UIMs close to Site 2.

The most intuitive explanation for such a change is that the ubiquitin moiety binds to both sites simultaneously and bridges them. It is easy to envision a di- or poly-ubiquitin species accomplishing this, but it is less obvious how a monomeric ubiquitin species (e.g. UbHis_6_) could do so. The ataxin3 tandem UIMs recognize the Ile-44 patch on the ubiquitin surface (46), and it has been suggested that Site 2 recognizes essentially the same region (34), which would preclude simultaneous binding. However, a more recent model suggests that Site 2 recognizes ubiquitin’s β1-β2 and α1-β3 loops (49), which would not overlap with the UIM binding site. Hence, the question of whether a single ubiquitin molecule can physically bridge Site 2 with the tandem UIMs remains open.

In conclusion, this work reveals new details of how ataxin3’s DUB activity is regulated, and hints at a complex interplay between the different domains of the enzyme and its substrates. Insights into how ubiquitin substrates can modulate allosteric changes in an enzyme like ataxin3 may prove relevant to other enzymes involved in the production and degradation of poly-ubiquitin chains.

## Methods

All proteins were heterologously expressed in *E. coli* and purified before use. Di-ubiquitin species were assembled *in vitro* using purified E1 & E2 enzymes. Cleavage of Ub-AMC was monitored in real time by measuring the increase in fluorescence at 445 nm. Detailed methods are provided in the Supplemental Information section, describing preparation of expression constructs, protein expression and purification, enzyme assays, and FRET experiments.

## Acknowledgements

This work was supported in part by grant R01NS065140 from the National Institutes of Health. The authors gratefully acknowledge expert advice from Marilyn Jorns and Shae Padrick and technical assistance from Hubza Syeda.

## Author contributions

MVR and PJL conceived and designed the research and wrote the paper. MVR performed the majority of the experiments, while PJL performed the FRET experiments. PJL and PDM prepared the FRET reporter construct. KCG prepared and purified many of the protein reagents used.

## Supporting Information

### Materials and Methods

#### Reagents

Enzymes for cloning and mutagenesis were purchased from New England Biolabs (Ipswich, MA). PCR primers were purchased from Integrated DNA Technologies Inc. (Coralville, IA) and sequencing of constructs was performed by Genewiz (South Plainfield, NJ). All chemicals were from Sigma-Aldrich (St. Louis, MO) unless otherwise stated, and media components were purchased from Fisher Scientific. Chromatography columns were obtained from GE Healthcare, and the Ub-AMC was purchased from Boston Biochem (Cambridge, MA).

#### Preparation of Expression Constructs

The preparation of expression vectors for full-length Q11 ataxin3 (corresponding to residues 1-345), the ataxin3 Josephin domain (residues 1-190), ubiquitin, and UbHis_6_ has previously been described (1, 2). Other expression constructs were prepared as described below. All PCR experiments used Phusion polymerase (New England Biolabs). Subcloning and mutagenesis primers are shown in Table S1. Cloning, mutagenesis and plasmid amplification were performed using the *E. coli* Mach1 strain (Invitrogen).

#### Ataxin3 single- and double-site mutants

Mutant constructs were prepared using a PCR-based mutagenesis protocol (3). The ataxin3 tandem UIM mutant was created in two sequential steps, first introducing the S236A codon change, followed by the S256A change. The “double-site mutant” (W87K plus S236A/S256A) was created by introducing the W87K mutation into the background of the tandem UIM mutant.

#### YPet-ataxin3-CyPet FRET construct

Plasmids encoding CyPet and YPet were obtained from Addgene (plasmids #14030 & 14031, respectively). The YPet and CyPet genes were PCR-amplified using the corresponding primers shown in Table S1, and the sequence corresponding to residues 1-265 of ataxin3 was amplified from the pETHSUL-ataxin3-Q11 template using the At3_FRET primer pair. The amplified products were purified using a QiaQuick PCR purification kit (Qiagen). The YPet sequence was then fused to the 5’ end of the ataxin3 gene using overlap extension. The fusion product was gel-purified, and the plasmid encoding the desired product was prepared by combining BseI-cleaved pETHSUL, the YPet-ataxin3 fusion product, and the amplified CyPet gene in a DNA assembly reaction using the NEBuilder HiFi DNA assembly kit (New England Biolabs). The resulting plasmid encodes a fusion protein containing His_6_-SUMO-YPet-ataxin3-CyPet, with a Gly_5_ linker between the YPet and ataxin3 sequences; residue 265 of ataxin3 is fused directly to the start methionine of the CyPet protein. After removal of the His_6_-SUMO tag, the mature fusion protein contains a single glycine residue at the N-terminus, immediately preceding the start methionine of the YPet protein.

#### Ypet

The YPet sequence was amplified from the Addgene plasmid #14031 using primers YPet_fwd and YPet-cntrl_rev. The resulting PCR product was purified with the QiaQuick kit and inserted into linearized pETHSUL using the NEBuilder HiFi DNA assembly kit.

#### Ataxin3-CyPet

The ataxin3 sequence was amplified from the pETHSUL-ataxin3-Q11 plasmid using primers At3-CyPet_fwd and At3_FRET_rev; the CyPet sequence was amplified from the Addgene plasmid using primers CyPet_fwd and new-CyPet_rev. After purification of the PCR products, the two inserts were combined with linearized pETHSUL using the NEBuilder HiFi DNA assembly kit.

#### MBP

The gene encoding MBP was PCR-amplified using primers MBP_pETHSUL_fwd and MBP_pETHSUL_rev, after which it was inserted into pETHSUL using ligation-independent cloning, following the published protocol (4).

#### MBP-UIM fusion

The sequence of MBP fused to a C-terminal Ala_5_ linker was amplified from pETXSH-MBP-5A (5), using primers MBP-5A_fwd and MBP-5A_rev.The sequence of the ataxin3 tandem UIM was amplified using primers UIM_fwd and UIM_rev. The MBP and UIM sequences were then fused using overlap-extension PCR and the fused amplicon was inserted into the BsaI and HindIII sites of the in-house vector pETHT; this vector encodes an N-terminal His_6_ tag and TEV-protease recognition site upstream of the protein being expressed. In a subsequent step, the sequence of the linker connecting MBP and the tandem UIM was changed from Ala_5_ to Gly_2_ using mutagenesis primers 5A-2G_mut_fwd and 5A-2G_mut_rev.

#### His_6_-MBP

An N-terminally His-tagged version of MBP was expressed from the in-house vector pETHM3c, which encodes residues 1-370 of MBP, with the sequences MKHHHHHHP and LEVLFQGP appended at the N- and C-termini, respectively.

#### Protein Expression and Purification

Ataxin3 full-length and Josephin-domain proteins were expressed and purified as described by Rao *et al.* (1); ubiquitin and UbHis_6_ proteins were expressed and purified as described by Weeks *et al.* (2). DiUb_48_ and diUb_63_ were prepared as described (2, 6). All purification steps were carried out at 4°C.

The YPet-ataxin3-CyPet, YPet, and ataxin3-CyPet proteins were prepared as follows. The appropriate expression plasmid was transformed into *E. coli* Rosetta (DE3) cells (Novagen). An overnight culture was diluted 20x into one liter of LB containing 100 µg/mL ampicillin. The culture was grown at 30°C to OD600 ∼0.8, induced with 0.5 mM IPTG, and incubated with shaking overnight at 24°C, after which the cells were harvested and frozen. Thawed cell pellets were suspended in Buffer A1 (50 mM Tris, pH 8.0, 250 mM NaCl, 10 mM imidazole, 0.1 mM TCEP) augmented with 2 µg/mL each RNase and DNase, 10 mM MgCl_2_, and protease inhibitor tablets (Thermo-Pierce, used per manufacturer’s instructions). In addition, the TCEP concentration in the suspension buffer was increased to 1 mM. Cells were lyzed by three passes through an Emulsiflex C5 cell disrupter (Avestin, Inc., Ottawa, Canada). The cell lysate was clarified by successive low- and high-speed centrifugations (15,300*xg* and 118,000*xg*, respectively) and loaded onto a 1-mL HiTrap IMAC-HP column (GE Healthcare) equilibrated with Buffer A1. After washing, the protein was eluted with Buffer B1 (50 mM Tris, pH 8.0, 250 mM NaCl, 250 mM imidazole, 0.1 mM TCEP). Protein-containing fractions were pooled; EDTA was added to a final concentration of 2 mM and 0.5 mg of purified dtUD1 SUMO hydrolase was added (4), after which the protein was dialyzed overnight versus 25 mM Tris, pH 8.0, 50 mM NaCl, 0.1 mM TCEP. Proteins were concentrated to roughly 2 mg/mL, flash-frozen, and stored at −80°C.

The MBP-UIM fusion protein and His_6_-tagged MBP were expressed in BL21(DE3) cells using self-inducing medium (7) for the former and IPTG induction (as described above for the FRET ataxin3 constructs) for the latter. Cells were grown at 30°C for 24 h, harvested, and frozen at −80°C. Upon thawing, cells were resuspended in Buffer A2 (50 mM sodium phosphate pH 7.4, 250 mM NaCl, 10 mM imidazole, 10% glycerol) plus 10 mM MgCl_2_ and 2 µg/mL each of DNase and RNase, and lysed by three passes through an Emulsiflex C5 cell disrupter. The cell lysate was clarified by successive low- and high-speed centrifugations at 17,600x*g* and 118,000x*g*, respectively. The clarified lysate was loaded onto an IMAC-HP column (GE Healthcare), which was washed with 20 column volumes of Buffer A2 and then eluted with Buffer B2 (50 mM sodium phosphate pH 7.4, 250 mM NaCl, 250 mM imidazole, 10% glycerol). Protein-containing fractions were combined, dialyzed against 25 mM Tris, pH 7.5, 50 mM NaCl, and loaded onto a HiTrap-Q HP column (GE Healthcare). Pure protein was eluted using a linear gradient from 50 mM to 1 M NaCl. The His_6_-tagged MBP was subjected to an additional size-exclusion purification step, using a HiPrep Sephacryl S-200 HR column equilibrated in 25 mM Tris, pH 7.5, 200 mM NaCl.

#### Enzyme Assays

Ub-AMC cleavage was monitored at room temperature (28 ± 1 °C) in a Tecan Spark plate reader equipped with a 410-nm dichroic mirror, using Corning black half-area plates (part no. 3993). For full-length ataxin3 constructs, the assay concentrations used were 20 nM DUB + 1 µM substrate. However, using these concentrations, the basal activity of the Josephin domain was too low to measure reliably, and thus for Josephin domain constructs, the assay was modified to use 50 nM DUB + 2 µM substrate. In all cases the reaction buffer was 20 mM Tris-Cl, pH 7.5, 5 mM DTT, with a final reaction volume of 100 µL. Sample was excited at 345 nm and fluorescence was read at 445 nm; a 10 nm bandwidth was used for both excitation and emission. The reaction was initiated by addition of enzyme, after which fluorescence was read at intervals of ca. 1 s for a duration of 5-10 minutes. Initial rates were calculated from linear fits of the fluorescence *vs*. time data. A set of standard concentrations of the AMC fluorophore (100-800 nM) was measured with each day’s experiments and was used to convert raw fluorescence data to molarity.

Control experiments were conducted in the absence of ataxin3 to ensure that no spurious fluorescence was contributed by the ubiquitin constructs (Fig. S2*A*). In addition, we observed no stimulation of DUB activity with the His_6_-tagged control protein His_6_-MBP (Fig. S2*B*).

#### FRET experiments

Fluorescent proteins were prepared at a concentration of 200 nM in 20 mM Tris-Cl, pH 8.0 + 5 mM DTT. Fluorescence measurements were made in 100-µL volumes in Corning black half-area plates (part no. 3993) using a Tecan Spark plate reader equipped with a 410-nm dichroic mirror. Samples were excited at 398 nm and the emission spectrum was read between 440 and 600 nm. A 5-nm bandwidth was used for both excitation and emission. Interacting proteins (ubiquitin constructs and control proteins) were added at a final concentration of 25 µM. The following control experiments were carried out (Fig. S3): *i)* The YPet-ataxin3-CyPet reporter construct was incubated with and without 25 µM SUMO (a ubiquitin-like protein that does not bind ataxin3); *ii)* An equimolar mixture of YPet and the ataxin3-CyPet fusion protein was examined with and without UbHis_6_ and diUb_48_ at 25 µM. No change in FRET was observed for either of these added proteins.

#### Modeling the fluorophore-fluorophore distance distribution in the YPet-ataxin3-CyPet reporter

A coarse-grained model was developed in order to define the range and distribution of possible donor-acceptor distances in the YPet-ataxin3-CyPet fusion protein. Starting with the YPet protein, we noted that the approximate distance from the fluorophore to the protein’s C-terminus (the site of attachment to ataxin3) is 23 Å. Given that the fluorescent protein is connected to the Josephin domain by a flexible Gly_5_ linker, this implies the YPet fluorophore can occupy a range of positions that sweep out a spherical surface with a radius of roughly 25 Å, centered about the Josephin N-terminus (yellow circle, Fig. S4*B*).

Between the C-terminus of the Josephin domain and the CyPet protein are 85 residues of ataxin3’s disordered C-terminal region (shown as a dashed gray line in Fig. S4*A-B*). Assuming this region behaves as a random coil, the mean end-to-end distance <*L*> can be approximated by <*L*^2^> = *L*_0_*N* (8). Using an estimate of 81.8 Å^2^ for *L*_0_ yields <*L*> ∼83 Å (9). Thus, the average position of the C-terminus of the ataxin3 protein should sweep out a sphere with a radius of 83 Å, centered about the Josephin C-terminus (gray circle, Fig. S4*B*). Obviously, the ataxin3 C-terminus can explore positions other than those defined by this average 83-Å radius, but limiting the calculation to the average radius is sufficient for the purpose of defining average donor-acceptor distances.

The CyPet protein is attached to the ataxin3 C-terminus; the distance from the CyPet fluorophore to the point of attachment is approximately 40 Å. Hence, the CyPet fluorophore is expected to sweep out a sphere of radius 40 Å (cyan circle, Fig. S4*B*), centered on the position of the ataxin3 C-terminus. Thus, in this coarse-grained model, the possible fluorophore-to-fluorophore distances are represented by all distances between those points on the surface of the cyan sphere and those points on the surface of the yellow sphere (with the understanding that cyan sphere can be centered at any position on the gray sphere).

Each sphere was approximated by a set of 74 points evenly distributed over its surface, giving rise to a total number of possible distances equal to 74^3^ = 405,224. A simple steric constraint was then applied that ignored any pair of YPet-CyPet points for which either point fell within a box that narrowly encompasses the Josephin domain; this reduced the number of pairwise distances to 315,962. These distances were found to be broadly distributed about an RMS value of 94.3 Å (Fig. S4*C*).

This simple geometric model predicts that, in the absence of any ubiquitin species, the separation between the YPet and CyPet fluorophores will vary widely, assuming that the ataxin3 C-terminal region behaves as a random coil. However, the RMS separation predicted by this model is substantially greater than the value of 54 Å inferred from the FRET data. This may imply that the ataxin3 C-terminal region, while intrinsically disordered, nevertheless adopts a persistence length shorter than that expected for a purely random coil (9). It is also worth noting that the YPet/CyPet pair exhibits a weak propensity for heterodimerization (10), which might tend to reduce the average fluorophore separation seen in the absence of ubiquitin.

**Table S1.**
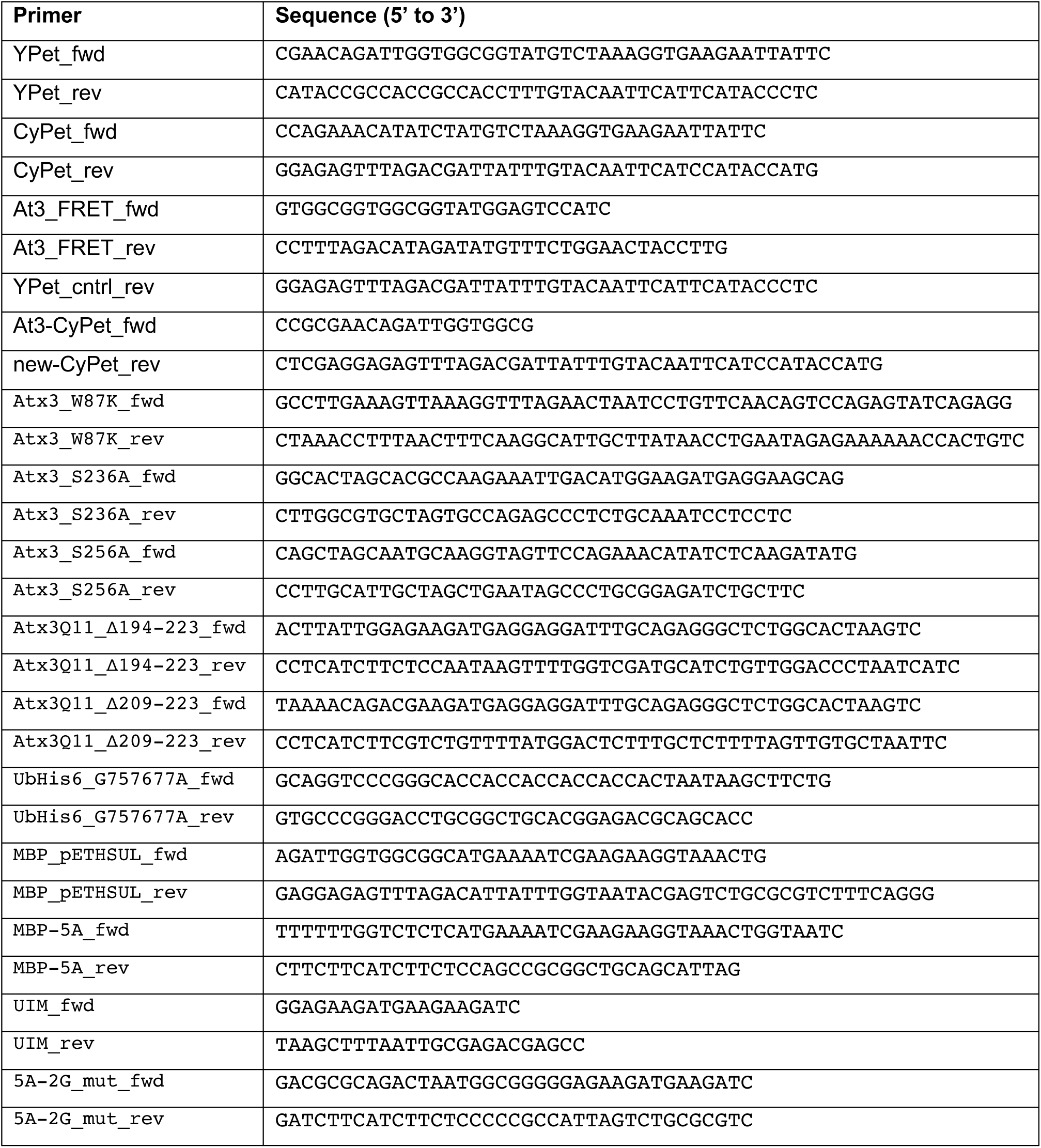
Primers used for construction of expression vectors.

**Table S2.** Basal activity rates for ataxin3 and Josephin domain.

| **Construct** | **Assay conditions** | **Basal activity**<br>**(mol product/mol enzyme per s)** |
| --- | --- | --- |
| Full-length | 20 nM enzyme/1 μM substrate | $4.9 \times 10^{-3} \pm 1.3 \times 10^{-3}$ |
| Josephin domain | 50 nM enzyme/2 μM substrate | $1.0 \times 10^{-3} \pm 0.3 \times 10^{-3}$ |

**Fig. S1.**
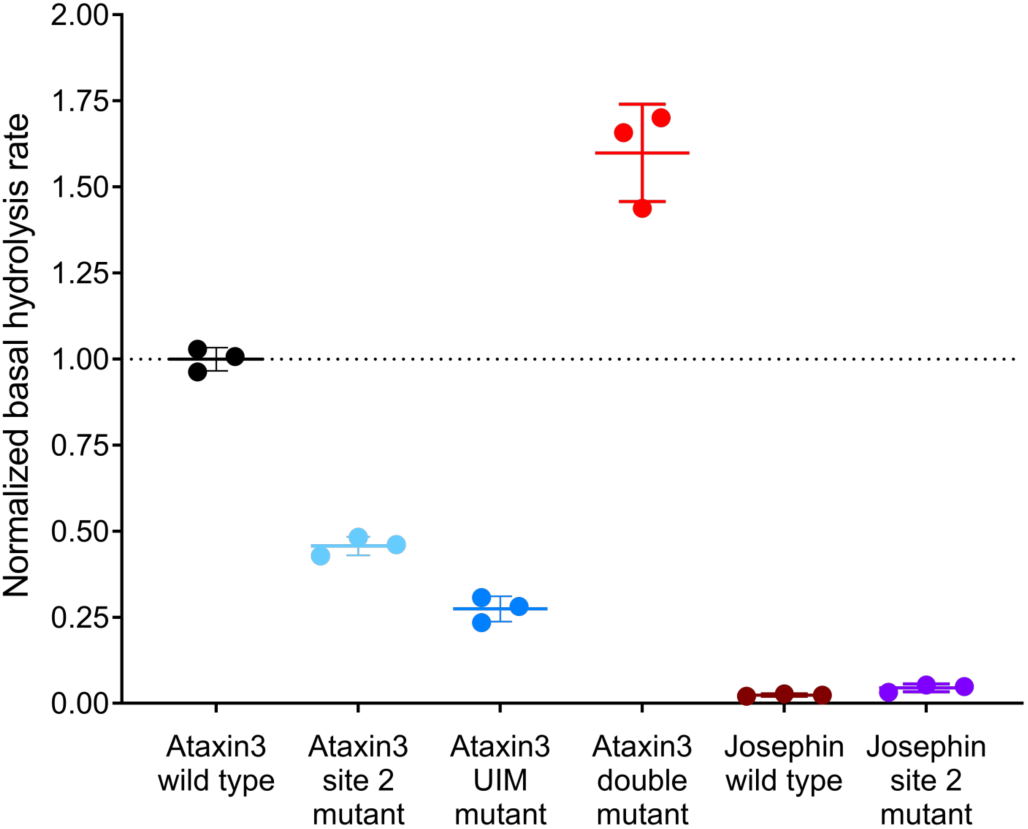
Comparison of basal activity levels for full-length ataxin3 and Josephin domain constructs. Basal activity rates for wild type and mutant constructs, calculated as the increase in fluorescence over time, and normalized to full-length ataxin3 wild-type activity. We were unable to reliably measure the basal activity for the Josephin domain under standard assay conditions; therefore, for comparison’s sake the experiments shown in this figure were performed with higher enzyme and substrate concentrations for all constructs tested (50 nM and 2 µM respectively).

**Fig. S2.**
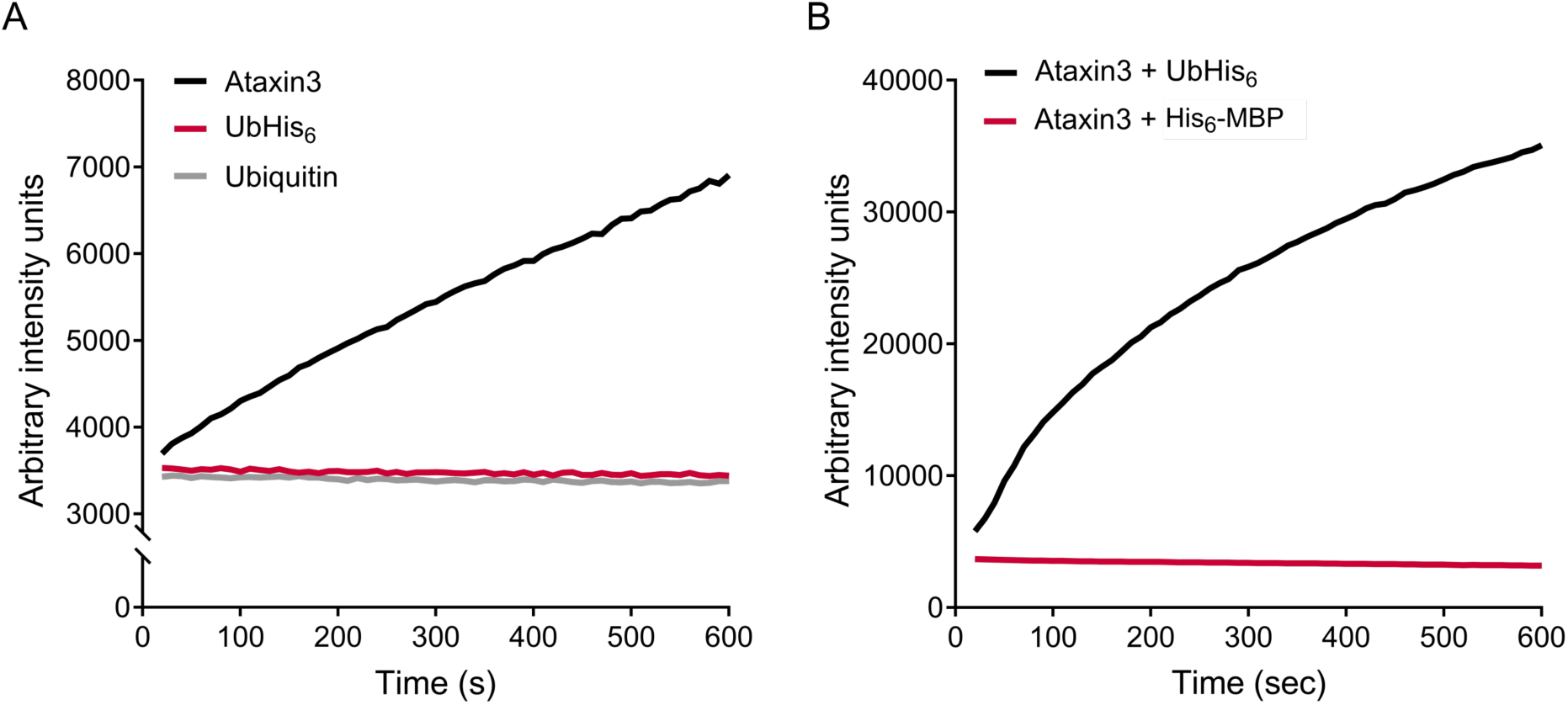
Enzyme assay control experiments. *(A)* Representative traces showing raw fluorescence counts over time for wild-type ataxin3 (black), and 75 µM UbHis_6_ (red) and ubiquitin (grey) in the absence of the DUB. No spurious fluorescence was detected for the ubiquitin constructs alone. *(B)* The control protein His_6_-MBP (75 µM) does not stimulate ataxin3 DUB activity. For all reactions, *n* ≥ 3.

**Fig. S3.**
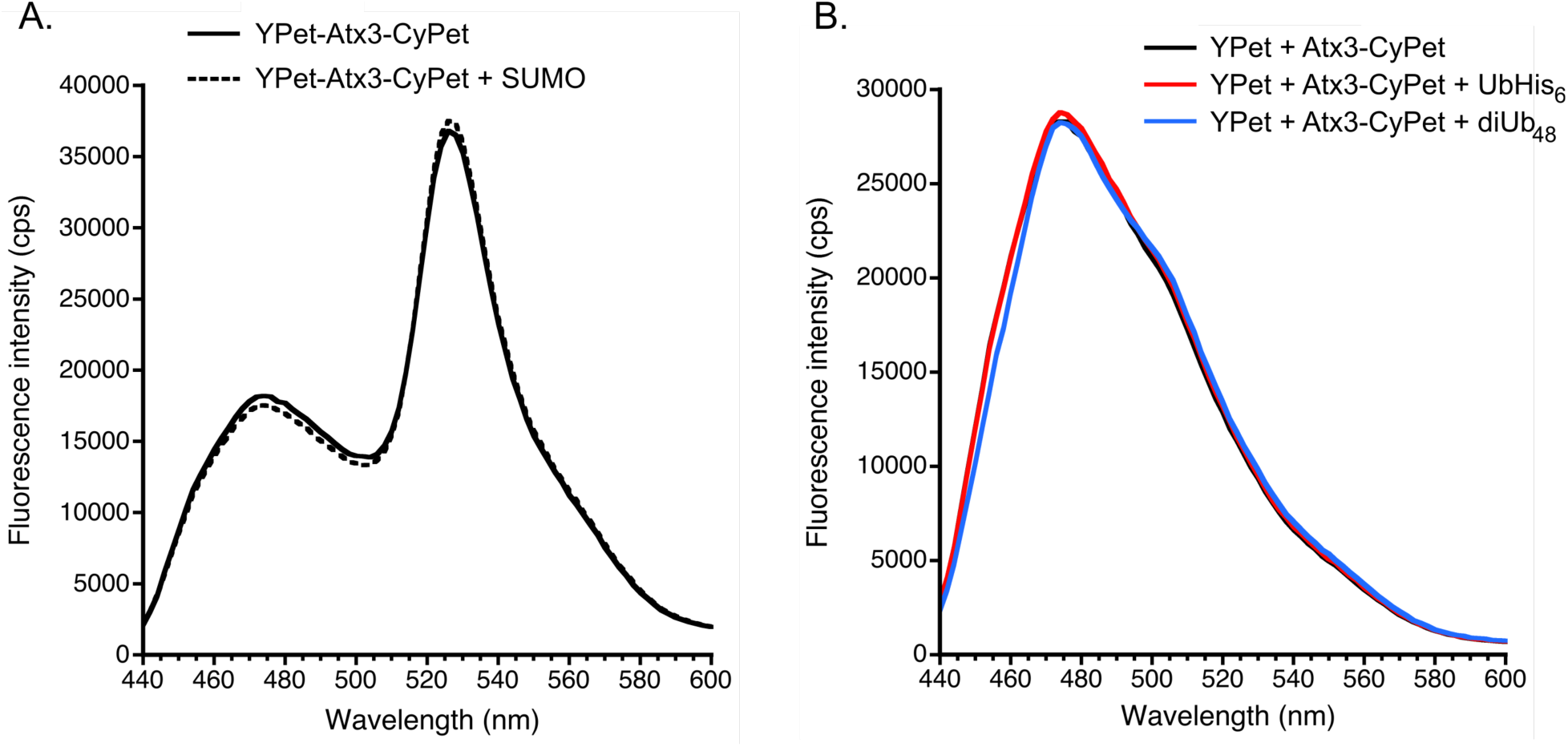
FRET control experiments. *(A)* FRET enhancement requires specific binding by a ubiquitin-containing construct. 25 µM concentrations of SUMO, a ubiquitin-like protein that does not bind ataxin3, do not elicit FRET. *(B)* Control experiments using equimolar quantities of YPet and ataxin3-CyPet ± 25 µM UbHis_6_ (red) and diUb_48_ (blue); no FRET is observed when the two fluorescent proteins are present *in trans*.

**Fig. S4.**
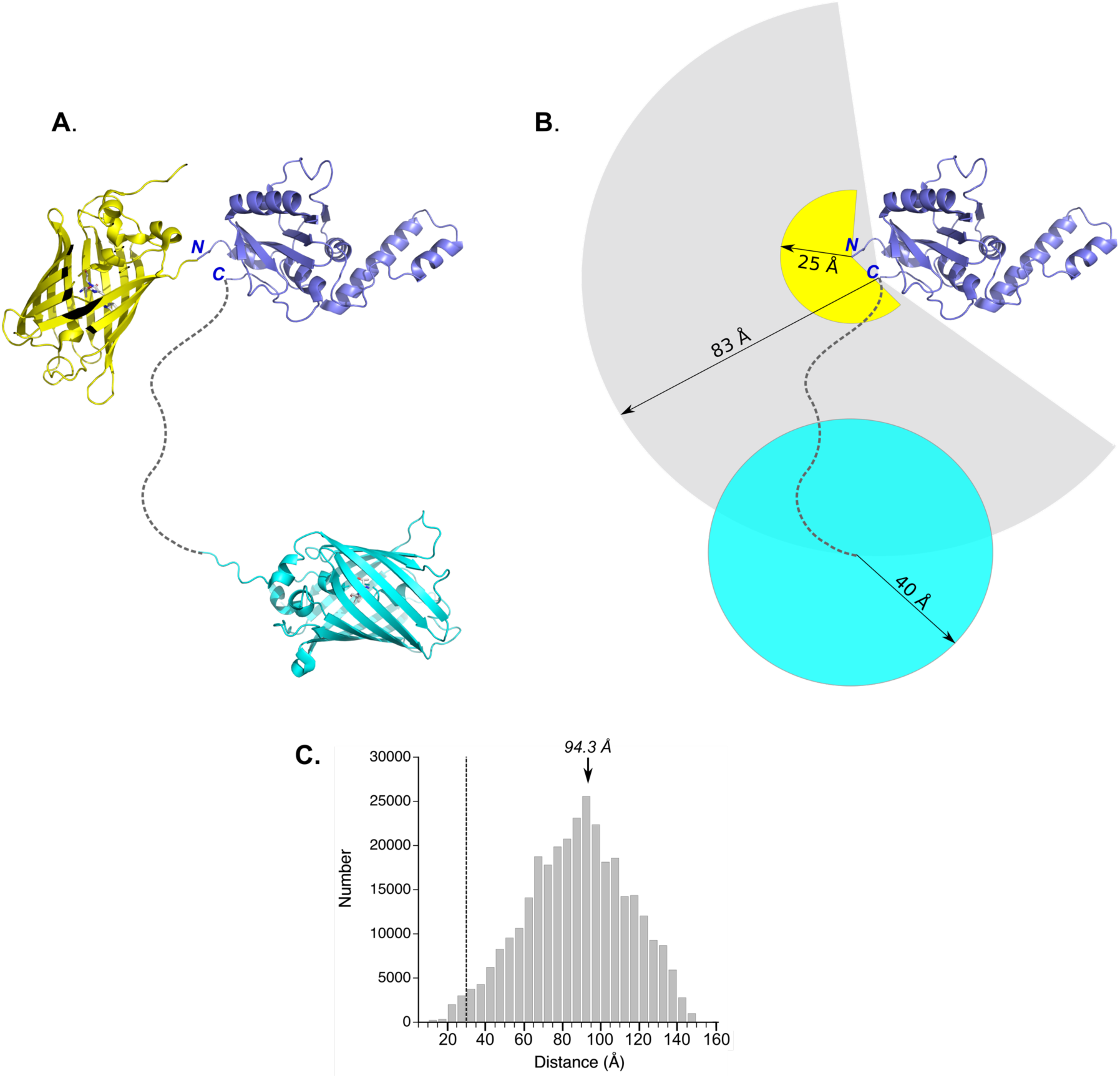
Coarse-grained modeling of the inter-fluorophore distances in the YPet-ataxin3-CyPet FRET reporter. ***(A)*** Schematic representation of the YPet-ataxin3-CyPet FRET reporter. Drawn to scale are YPet (yellow), the ataxin3 Josephin domain (blue), and CyPet (cyan). The disordered C-terminus of ataxin3 is represented as a dashed gray line. The N- and C-termini of the Josephin domain are indicated. Josephin and fluorescent protein coordinates were taken from PDB entries 1YZB & 5OXC, respectively (11, 12). ***(B)*** Simplified geometric model used to approximate the inter-fluorophore distance distribution in the YPet-ataxin3-CyPet reporter. The yellow circle is centered about the Josephin domain N-terminus, and represents the spherical surface swept out by the YPet fluorophore due to random motions of the protein. The gray circle is centered on the Josephin domain C-terminus, and represents the spherical area swept out by the disordered ataxin3 C-terminal region (represented by the dashed gray line), with an average end-to-end distance of 83 Å. The cyan circle represents the 40 Å sphere swept out by the CyPet fluorophore, and is centered at points on the surface of the gray sphere. ***(C)*** Distribution of possible fluorophore-fluorophore distances, as defined by the model described above. The RMS value of 94.3 Å is indicated by an arrow. The dashed line at a distance of 30 Å represents the lower limit for close approach, as determined by the protein shells surrounding the YPet and CyPet fluorophores (13).

